# Weighting of celestial and terrestrial cues in the monarch butterfly central complex

**DOI:** 10.1101/2022.01.25.477695

**Authors:** Tu Anh Thi Nguyen, M. Jerome Beetz, Christine Merlin, Keram Pfeiffer, Basil el Jundi

## Abstract

Monarch butterflies rely on external cues for orientation during their annual long-distance migration from Northern US and Canada to Central Mexico. These external cues can be celestial cues, such as the sun or polarized light, which are processed in the internal compass of the brain, termed the central complex (CX). Previous research typically focused on how individual simulated celestial cues are encoded in the butterfly’s CX. However, in nature, the butterflies perceive several celestial cues at the same time and need to integrate them to effectively use the compound of all cues for orientation. In addition, a recent behavioral study revealed that monarch butterflies can rely on terrestrial cues, such as the panoramic skyline, for orientation and use them in combination with the sun to maintain a directed flight course. How the CX encodes a combination of celestial and terrestrial cues and how they are weighted in the butterfly’s CX is still unknown. Here, we examined how input neurons of the CX, termed TL neurons, combine celestial and terrestrial information. While recording intracellularly from the neurons, we presented a sun stimulus and polarized light to the butterflies as well as a simulated sun and a panoramic scene simultaneously. Our results show that celestial cues are integrated linearly in these cells, while the combination of the sun and a panoramic skyline did not always follow a linear integration of action potential rates. Interestingly, while the weighting between the sun and polarized light was invariant between individual input neurons, it varied strongly when the sun stimulus and the panoramic skyline were presented simultaneously. Taken together, this dynamic weighting between celestial and terrestrial cues may allow the butterflies to flexibly set their cue preference during navigation.

## Introduction

Spatial orientation has been investigated behaviorally in many insects, ranging from desert ants (Wehner, 2003; Wehner and Müller, 2006), honeybees (Rossel and Wehner, 1984; Edrich et al., 1979; Brines and Gould, 1979), dung beetles (el Jundi et al., 2019; Dacke et al., 2021), and locusts (Homberg, 2015), to moths (Dreyer et al., 2018a; Dreyer et al., 2018b). This also includes the monarch butterfly *(Danaus plexippus)*, which covers a distance of about 4,000 kilometers on its annual migration to its overwintering spots in Central Mexico (Merlin and Liedvogel, 2019). During this long-distance migration, the butterflies use the sun as their main orientation reference (Stalleicken et al., 2005). To successfully maintain their southerly direction over the course of a day, the butterflies integrate time information from the antennae (Merlin et al., 2009; Guerra et al., 2012) and the brain (Sauman et al., 2005) into their sun compass. In addition to the sun, monarch butterflies may also rely on the polarization pattern of the sky for orientation (Reppert et al., 2004). While the pattern of polarized light is perceived by a specialized dorsal region of the monarch butterfly eye (Sauman et al., 2005; Stalleicken et al., 2006), the sun is detected by the main retina of the eye (Merlin et al., 2011). Skylight information is then transferred via the optic lobe and anterior optic tubercle to input neurons of the central complex (CX), termed tangential (TL) neurons (Heinze and Reppert, 2011; Nguyen et al., 2021). These neurons transfer skylight information from the bulb of the lateral complex to the central complex lower division in many insects (Held et al., 2015; el Jundi et al., 2018; Hensgen et al. 2020), including monarch butterflies (Fig. 1A). As shown for other insects (Stone et al., 2017; Hardcastle et al., 2021) TL cells synapse onto a network of heading-direction cells that flexibly encode the actual flight direction of an animal based on sensory-motor information (Seelig and Jayaraman, 2015; Turner-Evans et al., 2017; Green et al., 2017; Fisher et al., 2019; Kim et al., 2019; Okubo et al., 2020). While previous research focused on how the CX processes single celestial stimuli in the monarch butterfly brain (Heinze and Reppert, 2011; Nguyen et al., 2021; Beetz et al., 2021), the controlled lab conditions rarely reflected the cue conditions in nature. Thus, to obtain a highly robust compass network, multiple visual cues, such as the sun and polarized light are integrated simultaneously in nature (el Jundi et al., 2014; Lebhardt and Ronacher, 2014). Moreover, experiments on tethered flying monarch butterflies suggest that they combine a sun stimulus and a panoramic skyline to keep a directed flight heading (Franzke et al., 2020), similar to what has been reported for Australian bull ants (Reid et al., 2011) and honeybees (Towne and Moscrip, 2008; Towne et al., 2017). But how visual sceneries based on multiple stimuli, such as the sun and polarized light or the sun and a panoramic scene, are combined and how each of the cues is weighted neuronally has not been investigated in the monarch butterfly brain so far. To study this, we recorded intracellularly from TL cells in the monarch butterfly brain and analyzed how they respond to simultaneously presented cues, such as a simulated sun and polarized-light stimulus as well as a sun and panoramic-skyline stimulus.

**Fig. 1:**
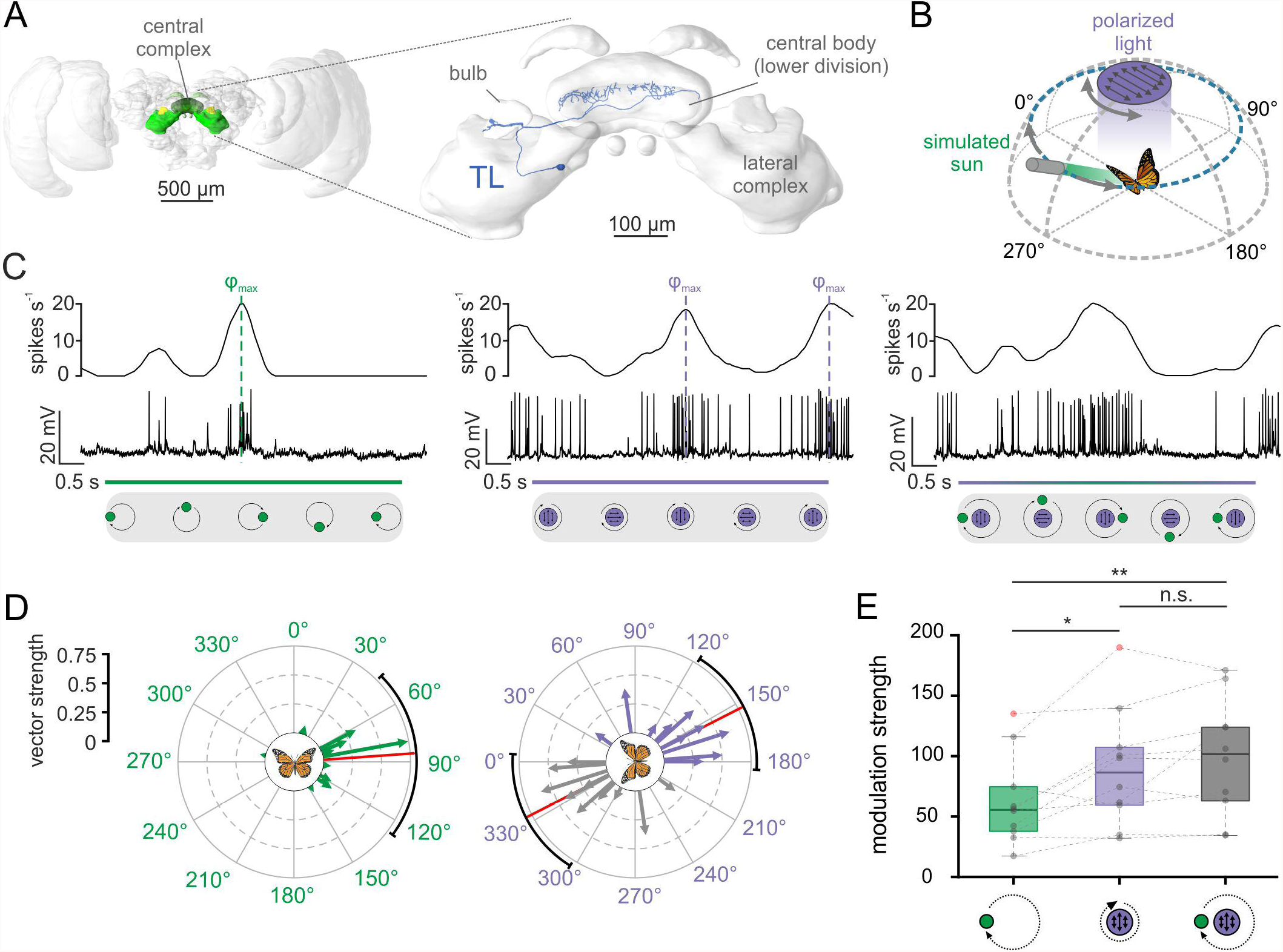
TL neurons of the central complex encode simulated celestial cues. **(A)** Left: Frontal view of the monarch butterfly brain. Highlighted in green are the neuropils of the central complex and lateral complex. Right: The central complex and lateral complex. A reconstructed tangential neuron (TL) is shown in blue. Modified from Heinze et al. (2013). **(B)** Schematic illustration of the presented simulated celestial cues. The polarization stimulus was positioned dorsally to the butterfly and was rotated by 360°. The angle of polarization was aligned with the antero-posterior axis of the animal at the beginning of the rotation. The sun stimulus (elevation: 30°) was moved on a circular path around the animal. The angle of polarized light was oriented perpendicular to the direction of the sun stimulus, simulating their spatial relationship in nature. **(C)** Neural tuning of the same TL neuron to a moving sun stimulus (left), a rotating polarizer (middle) and when both stimuli were presented simultaneously (right). The upper curve of each plot shows the sliding average of the action potential recordings (middle row). The lower grey boxes illustrate the position of the stimuli during a clockwise 360°-rotation indicated. Preferred firing directions (□_max_) to the sun (left) and polarized light (middle) are indicated by dashed vertical lines. **(D)** The preferred firing directions of the tested TL-neurons (n=10) in response to the sun stimulus (left) and the polarization stimulus (right). Each arrow represents a single neuron. Arrow length indicates the vector strength (directedness) of the neural tuning. The circular plots are labeled in relation to the animals’ body axis (see schematic at the plot’s center), with 0° being anterior to the animal. The mean preferred firing directions (sun stimulus: 85.88° ± 54.94°; polarized light: 152.96° ± 34.16°) are indicated by the red solid lines and the confidence intervals (95%) by the black arcs. **(E)** Modulation strength of neural activity (n=10 TL neurons) in response to the sun stimulus (left), polarized light (middle), and the combination of the sun and polarized-light stimulus (right). The neural modulation to the sun stimulus was significantly weaker than to polarized light p_GREEN vs. POL_ = 0.02, t = 2.99, n = 10; paired t-test) and the combination of the stimuli (p_GREEN vs. COMBO_ = 0.002, t = -4.38, n = 10; paired t-test), while the modulation strength to polarized light and the combined stimuli did not differ from each other (p_POL vs. COMBO_ = 0.23, t = -1.28; paired t-test). Grey circles show individual data points. Outliers are indicated in red. Dotted black lines connect data points that are obtained from the same TL-neuron. Boxes indicate interquartile range. Whiskers extend to the 2.5^th^ and 97.5^th^ percentiles. Black horizontal lines show the median. n.s.: not significant, *: p <0.05, **: p <0.01.

## Materials and Methods

### Animals

Adult monarch butterflies (*Danaus plexippus*) of both sexes were kept in an incubator (I-30VL, Percival Scientific, Perry, IA, USA) with a 12:12 hours light-dark cycle at 25°C in Würzburg (Germany). They were provided with 15% sugar solution *ad libitum*. Some animals were caught at College Station, TX, USA during their annual southward migration. These animals were kept in the incubator at an 11:13 hours light-dark cycle at 23°C during light and 12°C during dark phases. They were fed with 20% honey solution every second day.

### Preparation and electrophysiology

After clipping off wings and legs, the butterflies were attached to a custom-built holder using dental wax (Omnident, Rodgau Nieder-Roden, Germany). The head capsule was opened frontally and muscle and fat tissue above the brain were removed. At least one of the antennae remained intact to avoid a disruption of circadian inputs to the compass network (Guerra et al., 2012; Merlin et al., 2009). To access the central brain with the electrode, the neural sheath was removed using fine tweezers. Throughout preparation and subsequent neural recording, the brain was immersed in monarch ringer (150 mM NaCl, 3 mM KCl, 10 mM TES, 25 mM sucrose, 3 mM CaCl_2_).

To record intracellularly from individual TL neurons, micropipettes were drawn from borosilicate glass capillaries (inner diameter: 0.75 mm and outer diameter: 1.5 mm, Hilgenberg, Malsfeld, Germany) using a Flaming/Brown horizontal puller (P-97, Sutter Instrument Company, Novato, CA, USA). After loading the micropipette with 4% Neurobiotin (Vector Laboratories, Burlingame, UK, dissolved in 1 M KCl), it was filled with a 1 M KCl solution. The micropipette was connected to an electrode holder with a chloridized silver wire, which was attached to a micromanipulator (Leica Microsystems, Wetzlar, Germany). Another chloridized silver wire served as reference electrode and was inserted into the opened head capsule close the butterfly’s mouthparts. Detected signals were amplified 10x using a BA-03X bridge amplifier (npi Elelctronic GmbH, Tamm, Germany). The signal was digitized with sampling rates between 1-20 kHz using a digitizer (Power1401, Cambridge Electronic Design, Cambridge, UK). The neural activity was observed on a computer using the software Spike 2 (version 9.00, Cambridge Electronic Design).

### Celestial stimuli

To simulate celestial cues, the same stimulus was used as described in Nguyen et al. (2021). A rotation stage (DT-50, PI miCos GmbH, Karlsruhe, Germany) was dorsally positioned to the animals. For the polarized UV light stimulus, a polarizer was mounted on top of the rotation stage. Because monarch butterflies detect polarized light in the UV range (Stalleicken et al., 2005; Sauman et al., 2005) a UV-LED with an emission peak at 365 nm (LZ1-10UV00-0000, OSRAM Sylvania Inc., Wilmington, MA, US) and a quarter white diffuser (Nr. 251, LEE filters, Hampshire, UK) were placed behind a linear polarization filter (Bolder Vision Optik Inc., Boulder, CO, USA) in the center of the rotation stage. This allowed us to present equally illuminated polarized UV light to the butterflies. The sun stimulus was presented using an unpolarized green LED with an emission peak at 517 nm (LZ1-10G102-0000, OSRAM, Munich, Germany). This LED was mounted on one of four arms extending from the rotation stage. To control for the influence of wavelength information, a UV LED was attached to the arm opposite to the green LED. Both light spots were adjusted to an elevation of 30° relative to the animal’s head and provided unpolarized light. All light stimuli (unpolarized green/UV light, polarized UV light) were adjusted to a photon flux of about 1.4 × 10^14^ photons/cm^2^/s, measured with a spectrometer (Maya2000 Pro, Ocean Optics) at the position where the animal faced the stimuli during recordings. Since the recordings were obtained via two setups with identical equipment, the angular positioning of the stimulus varied slightly. The polarization stimulus had an angular extent between 9.6°-10.4° at the butterfly’s eye. The angular size of the unpolarized light spots was 1.3-1.4°. The movements of the rotation stage were controlled via a custom-written script for the software MATLAB (Version R2019b, MathWorks, Natick, MA, USA). During the experiments, the rotation stage was turned by 360° in clock- and counterclockwise direction at a constant velocity of 60°/s while testing the response of the TL cells to a single stimulus (sun stimulus *or* polarized light) or a combination of the stimuli (sun stimulus *and* polarized light). As five of the TL neurons were obtained from migratory butterflies, we first tested if they differed in their general tuning characteristics from the recordings obtained in non-migratory butterflies. However, we did not find any differences in the relevant response characteristics tested here between both groups and decided to pool the data.

### Panoramic skyline and sun stimuli

The panoramic skyline was simulated via an LED arena consisting of a circular array of 128×16 RGB-LEDs (M160256CA3SA1, iPixel LED Light Co., Ltd, Baoan Shenzhen, China). The arena covered a visual field of 360° along the horizontal and 43° along the vertical plane around the animal. The LEDs were controlled via a Raspberry Pi (Model 3B, Raspberry Pi Foundation, Cambridge, UK). We presented the same panoramic skyline to the butterflies that has been used in recent behavioral experiments on monarch butterflies (Franzke et al., 2020). Each LED above the horizon was adjusted to a photon flux of about 6.68 × 10^10^ photons/cm^2^/s in the blue range (emission peak: 458 nm). LEDs below the horizon were turned off. The panoramic scene was uploaded as an RGB image 8bits/channel) to the Raspberry Pi and a custom-written program written in Go controlled the rotation movements of the stimulus. To avoid any dark adaption of the animals’ eyes, a panorama with a flat horizon (*flat panorama*) was presented to the animals throughout the entire experiment, while the panoramic scenery with the variable height profile (*panoramic skyline*) was used as a test stimulus. Intensity differences between the two panoramic stimuli were minimized by turning on a similar number of LEDs in both panoramas. To find TL neurons during our experiments, we first stimulated the animal with zenithal polarized light. Once a neuron responded to polarized light, the polarization stimulus was turned off and the panoramic skyline was presented and rotated by 360° around the animal in clock- and counterclockwise direction (at a constant velocity of 60°/s). To combine the panoramic scene with a sun stimulus, one LED above the horizon at an angular elevation of 18.9° was switched to a wavelength emission peak of about 516 nm and an intensity of about 6.14 × 10^12^ photons/cm^2^/s. We combined the sun stimulus either with the flat panorama or with the panoramic skyline. For the former, we moved the sun stimulus around the animal, while the flat panorama stayed stationary. For the latter, we rotated both stimuli around the animal.

### Histology and imaging

To evaluate anatomically the neuron type from which we recorded, Neurobiotin was iontophoretically injected into the cells (1-3.5 nA) for 3-5 min at the end of each experiment. After allowing the Neurobiotin to distribute for 20 min, the brains were dissected out of the head capsule and fixated for 18-24 h at 4°C in a sodium-phosphate buffer containing 4% paraformaldehyde, 0.2% picric acid, and 0.25% glutaraldehyde. They were then rinsed 4 × 15 min in 0.1 M phosphate buffered saline (PBS) and, afterwards, incubated with either Cy3-conjugated to streptavidin (Thermo Fisher Scientific, Waltham, MA USA, 1:1000) or Alexa568-conjugated to streptavidin (Molecular Probes, Eugene, OR, USA, 1:1000) diluted in PBS containing 0.3% Triton-X 100 (PBT) for 3 days at 4°C. The brains were then rinsed with PBT (3 × 20min) and afterwards with PBS (2 × 20min), before they were dehydrated through an ascending ethanol series (30, 50, 70, 90, 95, and 100%; 15 min each). Afterwards, the brains were immersed in a 1:1-mixture of ethanol and methyl salicylate (Fisher Scientific GmbH, Schwerte, Germany) for 20 min and then in 100% methyl salicylate for about 1 h at room temperature. The brains were then mounted in Permount (Fisher Scientific) between two cover slips with ten reinforcement rings (Avery, Toronto, Canada) as spacers. Finally, they were imaged using a confocal microscope (Leica TCS SP8, Wetzlar, Germany) with a 10x air objective (HCX PL-Apo 10x/0.4 CS, Leica).

### Data analysis

To consider a neuron for analysis, the following criteria had to be fulfilled: *i*. stable baseline during stimulus presentation, *ii*. spike amplitudes clearly above noise level and *iii*. distinct immunolabeling of the recorded neuron. If a neuron passed these criteria, the recorded file was imported into MATLAB for further analysis via custom-written scripts that included the CircStat toolbox (Berens, 2009). Events during stimulation were detected based on a manually set threshold and were assigned to a particular polarization angle during the polarizer rotation or to a corresponding azimuthal angle during circling of a light spot. Neuronal spiking rates were estimated by low-pass filtering the instantaneous firing rate of the action potentials and illustrated as sliding averages (Gaussian filtered, window size: 0.5 s) in the results. The preferred angles in response to the stimuli were determined as the mean vector of the bimodal (polarized light) or unimodal (sun stimulus) distribution of stimulus angles at the times of action potentials. The degree, to which the action potentials were clustered at this angular position, was defined as the vector strength. This value ranges from 0 to 1, with higher values indicating more directed responses

In addition to the aforementioned parameters, responses of each trial were binned into 18 bins and spike rate in each bin was calculated. This was used to obtain the modulation strength as described by Labhart (1996) using the following equation:

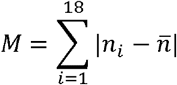

where *n* is the spiking rate (in spikes/s) and 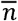 is the average spiking rate over the whole stimulation period. The higher the modulation strength, the stronger is the response of a neuron to a certain stimulus.

To predict the neural response to a combination of stimuli and to define the relevance of each cue on the neural coding, a weighted linear model was applied. This was based on the responses of the same neuron to the individual stimuli using the following equation:

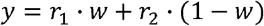

where *r*_*1*_ and *r*_*2*_ describe the actual response of a neuron to an individual stimulus (e.g., sun stimulus and polarized light for the combined celestial stimuli condition), respectively and *w* indicates the weighting of them. If *w* < 0.5, the response of *r*_*2*_ is weighted higher, while *w* < 0.5 indicates that the neural response of *r*_*1*_ dominates the response. To identify the weighting that matches best the actual neural response to the combination of both stimuli, we calculated the correlation between the actual neural response and the modeled neural responses based on different linear weightings. The weighting that exhibited the highest correlation coefficient, was then considered for further analysis.

As we did not have any prediction of whether and how the TL neurons respond to the panoramic skyline, we used the inter-trial difference between single stimulus presentations as reference. Inter-trial differences were determined by treating clock- and counterclockwise responses separately. At first neural responses were grouped based on the stimulus rotation. Then, for both rotation directions, we calculated the mean neural response and correlated them with the response from each trial. Finally, the correlation coefficients were averaged within and then across rotation groups. The closer the correlation were to unity, the more similar were the neural responses across trials. To quantify the neural response to the panoramic skyline further, the averaged neural response for clock- and counterclockwise rotations was compared with the response to the flat panorama during a 6 s time frame. Both correlation values were averaged across rotation groups again. If the neurons encoded parts of the panoramic skyline, we expected that the neural response to the panoramic skyline and the response to the flat panorama correlate less with each other than the neural responses across trials.

To further characterize the response of the TL cells to the panoramic skyline, we applied an intensity-based model for different elevations. Circular, excitatory TL neurons’ receptive fields were modeled by a Gaussian Kernel (11.5° width ± 2.9° standard deviation, corresponding to 5 pixels width and 2 pixels standard deviation of the arena) in MATLAB. This receptive field size is in a similar range as the excitatory component of measured locust TL neuron receptive fields (Takahashi et al. 2022). When the scene is rotated around the animal, this evokes changes in brightness in the modeled receptive field that are correlated with the silhouette of the panorama. Thus, if a bright sector of the panorama moves through the receptive field, it increases the spiking activity. In turn, if a dark sector of the panorama is moved through the receptive field, it will decrease the spiking activity. However, this change in spiking activity depends on the elevation of the receptive fields (Fig. 3E), which may vary in monarch butterfly TL neurons, as shown for the homologous neurons in fruit flies (Seelig and Jayaraman, 2013). We therefore defined each row of the LED arena between -16.1° and 16.1° as the center of a receptive field and modeled curves of the predicted neural modulation at different elevations when the panorama was rotated. To find the best match between the modeled response and the recorded neural response at different elevations, we calculated for the modeled responses at each elevation the cross correlation with the measured neuronal response. The modeled curve that exhibited the best match to the recorded neural response curve was included for further analysis.

### Statistics

To identify whether action potentials in response to the simulated skylight cues are non-uniformly distributed, we applied the Rayleigh-test (significance level <0.05). To test whether the preferred directions are significantly clustered around the 0°-180° axis (polarized light) and around 90° (sun stimulus), we used the V-test (significance level <0.05). To test for normal distribution and similar variances of the modulation strengths, the Shapiro-Wilk test and the Levene-test were employed, respectively. If data were normally distributed and exhibited the same variance, parametric hypothesis tests were applied (unpaired t-test and paired t-test, respectively). Otherwise, non-parametric tests were used (Wilcoxon-rank-sum-test for unpaired and the Wilcoxon signed rank test for paired mean values). For partially paired data, like the observed weighting factors, a mixed linear model was used to test if the mean values differed significantly between the two test groups. Averaged parameters are shown as mean ± standard deviation if not mentioned otherwise.

## Results

To understand how the monarch butterfly compass integrates multiple visual stimuli, we presented different visual stimuli in isolation and in combination to the animals while recording intracellularly from TL neurons (in fruit flies termed ring neurons) of the central complex (Fig. 1A). We successfully obtained recordings from 34 TL-neurons. 15 TL-neurons were tested with a combination of different celestial cues and 19 TL-neurons with a combination of celestial and terrestrial wide field stimuli. Of the latter group, all TL cells were tested with the panoramic skyline, 15 of them were exposed long enough to the flat panorama to be included in the inter-trial response analyses and 13 of them were presented the combined sun stimulus and panoramic skyline.

### Celestial cue integration in TL neurons

We first tested the neural tuning to simulated celestial cues. Similar to previous experiments (Heinze and Reppert, 2011; Nguyen et al., 2021; Beetz et al., 2021), a moving green light spot served as a sun stimulus while a rotating polarizer illuminated by UV light from the zenith was used to examine polarization sensitivity (Fig. 1B). To simulate the natural spatial relationship between the sun and polarized light, we oriented the polarization angle perpendicular to the sun-stimulus direction (Fig. 1B). As demonstrated earlier (Heinze and Reppert, 2011; Nguyen et al., 2021), TL neurons responded to both the sun stimulus and polarized light (Fig. 1C, left and middle graph). Interestingly, the highest action potential rates (preferred directions, □_max_) of the TL neuron to the sun stimulus and the polarization stimulus matched the 90°-relationship of the cues in nature. To investigate if this was true for all recorded TL cells, we analyzed the spatial distribution of the preferred directions (□_max_) in response to the sun stimulus and the polarization stimulus. The preferred directions to the sun stimulus were clustered around 90° in response to the sun stimulus (p = 0.008, v = 5.43; V-test; n = 10; Fig. 1D, left) and along the 0° - 180° axis in response to polarized light (p = 0.03, v = 4.37; V-test; n = 10; V-test; Fig. 1D, right). Taken together, the spatial relationship between the mean preferred directions of the recorded TL neurons when presenting sun and polarization stimulus in isolation matched the natural spatial relationship between both celestial cues.

When we presented both stimuli simultaneously, the neural tuning resembled a mixed response (Fig. 1C, right graph), suggesting that TL neurons integrate both stimuli in a weighted manner. To quantify which of the two stimuli dominated the neural response, we compared the modulation strengths in response to the single stimuli (sun stimulus *or* polarized light) with the modulation strengths of the same neurons in response to the combined stimulus (sun stimulus *and* polarized light). The modulation strength in response to the sun stimulus (62.48 ± 37.05, n = 10) was significantly weaker than the modulation strength of the same neurons to the polarization stimulus (89.81 ± 48.64, n = 10; p = 0.02, t =2.99; paired t-test) and to the combination of the stimuli (98.90 ± 48.43, n = 10; p = 0.002, t =-4.38; paired t-test; Fig. 1E). The modulation strength did not differ between the response to the polarizer and to the combination of the stimuli (p = 0.23, t =-1.28; paired t-test). This indicates that the response to the combined stimuli is shaped by the polarization input which seems to be weighted stronger than the sun-stimulus input.

### Weighting of celestial cues in TL neurons

To examine whether polarized light truly dominates the response to the celestial cues and how they are weighted in TL neurons, we next combined the responses to the isolated stimuli (Fig. 2A, upper plot) in a weighted linear model and calculated a predicted response to the combined stimuli. By varying the weight between the neural response to the polarizer and sun stimulus, we modeled different expected neural responses to the combined stimuli and correlated these modeled responses with the actual response to the combined stimuli (Fig. 2A, blue curve of lower plot). The modeled response with the highest similarity to the actual response of the neuron (Fig. 2A, red curve of lower plot) was selected to determine the neuron-specific cue weighting that indicates which cue dominated the neural response (Fig. 2A, lower plot). A weighting factor of 0 indicated that the combined response was entirely characterized by the sun stimulus while a weighting of 1 represented a tuning that was purely described by the polarization input.

**Fig. 2:**
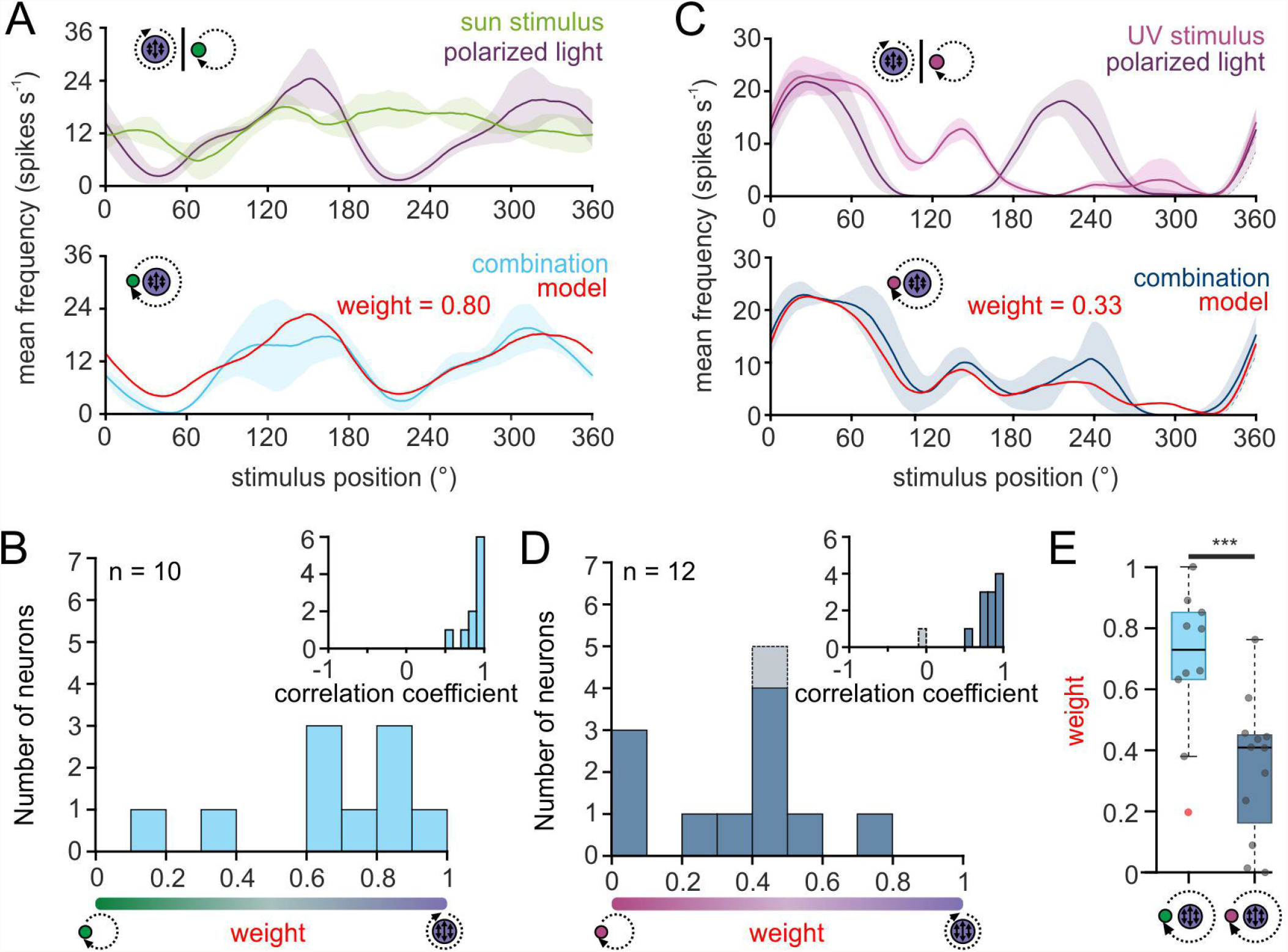
Weighting of celestial cues in TL neurons. **(A,C)** Upper plot: sliding averages of the action potential rates of two TL neurons to the green sun stimulus (A, green curve) or UV light spot (C, magenta) as well as to the polarization stimulus (A,C, violet curves) are shown. Lower plot: The response of the same neurons as in A, C when both stimuli were presented simultaneously (A, light blue; C, dark blue curve). Based on the response to the single cues, a weighted linear model was fitted to the data (red curve) and a weighting of the single cues was calculated. A weight between 0 and 0.5 suggests that the sun stimulus (A) or the UV light spot (C) dominated the combined response while a weight between 0.5 and 1 indicates that the polarization input dominates the combined response. The shaded areas show the standard deviation. **(B**,**D)** The distribution of the weighting factors for experiments with the sun stimulus and polarized light (B, n=10) and the UV light spot and polarized light (D, n = 12). Insets in B and D: The correlation coefficient shows how well the weighted linear model matched with the actual response to the combined stimulus. Weighting factors that were obtained from a low correlation coefficient, indicating that the neural response did not follow the weighted linear model, are shown in grey. **(E)** Comparison between the weighting factors obtained for the experiments with the green sun stimulus and polarized light (left) and the UV light spot and polarized light (right). The data are the same as in B,D. While the weighting is shifted to the polarization input when a green sun stimulus was combined with polarized light, the weighting is significantly shifted in favor of the light spot, when a UV light cue was combined with polarized light (p <0.001, F = 113.31; linear mixed model ANOVA). Grey circles show individual data points and red circles show outliers. Boxes indicate interquartile range. Whiskers extend to the 2.5^th^ and 97.5^th^ percentiles. Black horizontal lines show the median. ***: p < 0.001.

For all neurons, the correlation coefficients obtained through the comparison of the modeled and actual neural response were relatively high (Fig. 2B, inset, 0.86 ± 0.12, n = 10), suggesting that the response of TL neurons to the combination of celestial stimuli can be well described by a linear model. When presenting sun stimulus and polarized light simultaneously, most TL cells responded stronger to the polarization information (Fig. 2B), although the light intensity of the stimuli was set to the same photon flux. The bias towards the polarization stimulus could be caused by differences in the spectral content of the stimuli which may induce a difference in the relative brightness detected by the monarch butterfly eyes. Thus, we assumed that the UV polarization stimulus may appear brighter to the butterflies than the green sun stimulus due to a higher sensitivity of the photoreceptors to UV light. To test whether this may explain the dominance of the polarization stimulus on the neural response to the combined stimuli, we repeated the experiments with a UV sun stimulus that had the same photon flux as the polarization stimulus (Fig. 2C). Again, except for one neuron, the weighted linear model described the actual neural response well (Fig. 2D, inset, 0.77 ± 0.28, n = 12), which further confirms that celestial information is linearly integrated in TL neurons. In contrast to the trials with the green sun stimulus and polarized light, the weighting to the combined UV stimuli shifted in favor of the unpolarized UV-stimulus (Fig. 2D). The observed weightings differed significantly between the experiments with the green sun stimulus and polarized light (0.65 ± 0.28, n = 10) and the UV sun stimulus and polarized light (0.30 ± 0.23, n = 12; p < 0.001, F = 113.31; ANOVA, Fig. 2E). Taken together, our data show that TL neurons combine celestial cues linearly in monarch butterflies. However, the weighting between the sun and polarization input is highly affected by the spectral content and relative brightness of the presented stimuli.

### TL neurons are tuned to a panoramic skyline

In addition to celestial cues, recent experiments in *Drosophila melanogaster* suggest that central-complex neurons encode the entire visual scenery around the animal (Seelig and Jayaraman, 2015; Kim et al., 2019). One salient cue in a visual scene that can be used by many insects for orientation is the profile of a panoramic skyline (Graham and Cheng, 2009; Reid et al., 2011; Legge et al., 2014; Franzke et al., 2020). To understand how the monarch butterfly central complex encodes the panoramic skyline, we placed the butterflies at the center of an LED arena and recorded the neural activity of TL neurons while the animals were exposed to a panoramic skyline that was presented at the inner surface of the LED arena. We used the same panoramic skyline that has recently been used to study the monarch butterfly orientation behavior (Franzke et al., 2020; Fig. 3A). When we rotated the scene around the butterflies, we found that many TL neurons were modulated by the panoramic stimulus (Fig. 3B). To determine whether or not these modulations were spontaneous modulations of the neurons’ spiking activity, we analyzed the inter-trial variability of the neural activity (Fig. 3B, upper panel) and correlated the neural activity in response to each trial with the averaged response. As a control, the neural activity of the TL neurons in response to a flat panorama (Fig. 3B, lower panel) was correlated to the averaged response to the panoramic skyline. Neural responses to the panoramic skyline of the neurons across trials were highly correlated with their averaged response tuning (Fig. 3C, upper plot) indicating a low inter-trial variability. In contrast, the neural response to the flat panorama was poorly correlated with the averaged response to the panoramic skyline (Fig. 3C, lower plot). The correlation coefficients between inter-trial stimulus presentations of the panoramic stimuli were significantly closer to the averaged response tuning to the panoramic skyline (0.76 ± 0.07) than to the responses to the flat panorama responses and the averaged responses to the panoramic skyline (−0.02 ± 0.29; p < 0.001, sign rank = 120, Wilcoxon signed rank test, Fig. 3D). This demonstrates that monarch butterfly TL neurons encode, in addition to skylight cues, panoramic skylines.

**Fig. 3:**
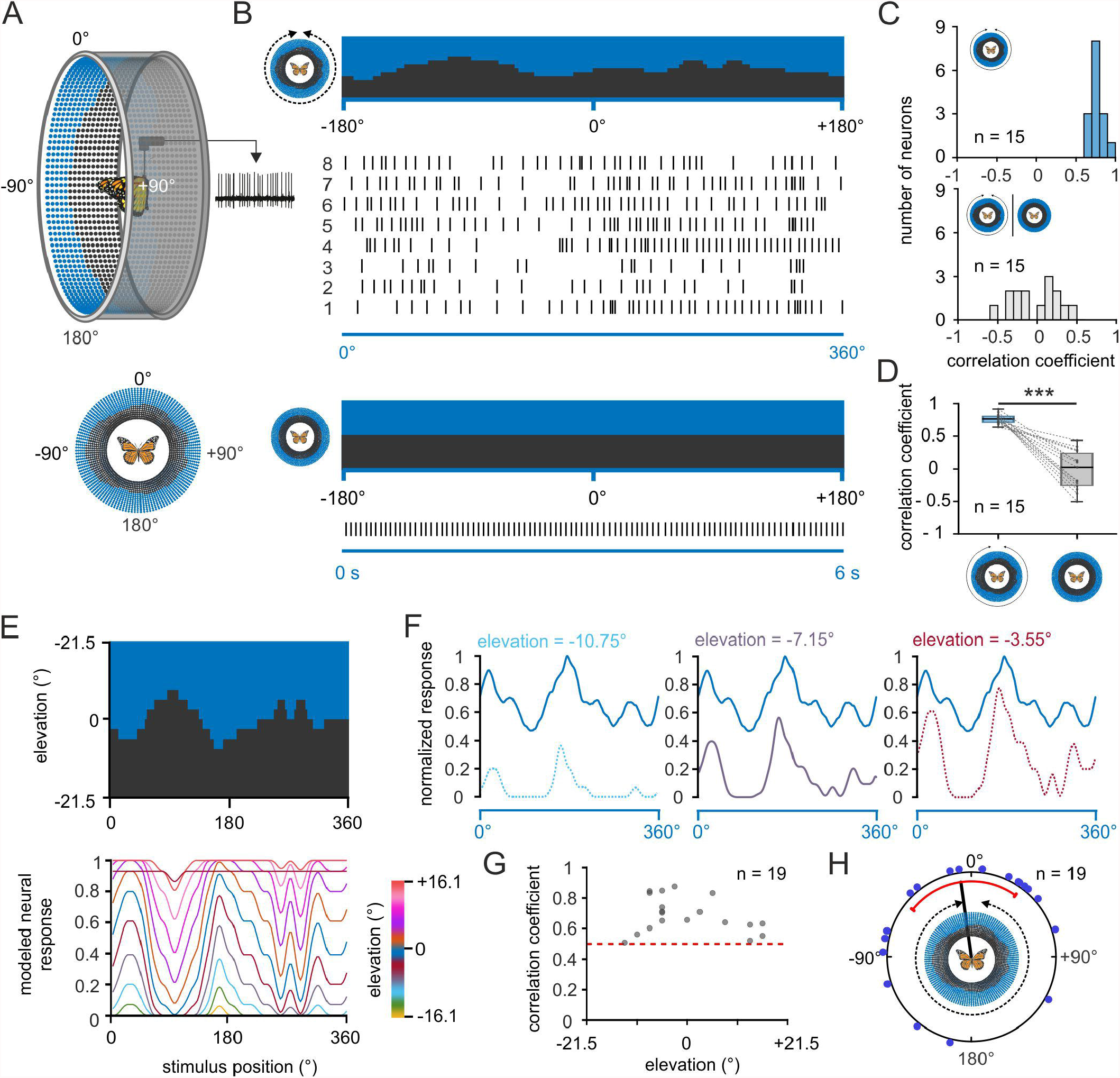
Response of central-complex TL neurons to a panoramic skyline. **(A)** Schematic drawing of the LED arena (top) that was used to test the coding of a panoramic scene in TL neurons. 0° was defined as the direction anterior to the butterfly, before the panoramic scene was rotated by 360° around the animal (bottom). **(B)** The response of a TL neuron to eight 360°-rotations of the panoramic scene is shown as raster plot (middle). Each vertical line represents an action potential. The orientation of the panoramic scene prior to rotation is shown (top). The TL neuron does not spontaneously modulate its action potential rate when a flat panorama is presented for 6s (below). **(C)** Distribution of correlation coefficients when comparing the neural response from each trial with the averaged response (upper panel) and when comparing the neural response to the flat panorama with the averaged response to the panoramic skyline (lower panel). **(D)** Comparison of the correlation coefficients obtained in C (p < 0.001; sign rank = 120, Wilcoxon signed rank test). Paired data points across the tested groups are connected by dashed lines. Boxes indicate interquartile range. Whiskers extend to the 2.5^th^ and 97.5^th^ percentiles. Black horizontal lines show the median. ***: p < 0.001. **(E)** Predicted neuronal response curves to the panoramic scenes of TL neurons that have visual fields centered at different elevations (middle). **(F)** The experimentally observed neural response of one recorded TL neuron (blue curve) to the panoramic scene plotted against three modeled changes in spiking activity of cells that detect light at different elevations (see E). The actual recorded response showed the best match to the modeled modulation at an elevation of -7.15 (middle plot; correlation coefficient = 0.84). **(G)** Predicted elevations of the visual fields for TL neuron response (grey dots; n=19) plotted against the corresponding correlation coefficient. Red dashed line indicates a correlation coefficient of 0.5. **(H)** Azimuthal shifts leading to the maximum correlation between the neural response and the panoramic skyline are plotted for each neuron (blue dots; n = 19). Shifts were biased towards the frontal visual field (p = 0.045; Z = 3.06, n = 19, Rayleigh test). The mean is indicated by a black solid line and the confidence intervals (95%) by a red arc.

As the receptive fields of the TL neurons can cover patches of different elevation and azimuth (Seelig and Jayaraman, 2013; Takahashi et al. 2022), it is not trivial to predict the neural response to the presented panoramic skyline. We noticed that the modulation of the spiking activity of the TL neurons seemed to correlate negatively with the troughs of the profile (Fig. 3B), suggesting that the cells are tuned to changes in brightness during the stimulus rotation. We therefore next modeled the change in the neural response for different receptive-field elevations based on the assumption that neurons encode changes in brightness associated with the panoramic skyline (Figs. 3E,F). For each of the modeled response, we calculated its cross correlations with the measured neuronal response. Although the receptive fields of the TL neurons were centered at all possible elevations, our model indicated that most cells had a receptive field centered at elevations between -10.75° and +10.75° (Fig. 3G), the segment outlining the panoramic skyline’s horizon. In addition, cross correlations also revealed which panorama azimuthal position gave the strongest neuronal response. The shifting distance between the observed TL response and the initial panorama position clustered in the anterior field of the animal (Fig. 3H; (p = 0.045; Z = 3.06, n = 19, Rayleigh test).

### Weighting of the sun and panoramic scene in TL neurons

We next wondered how TL neurons encode a visual scene that included both a simulated sun and the panoramic scene. As demonstrated in the example TL neuron, the moving sun stimulus mainly dominated the neural response, irrespective of the absence/presence of the panoramic skyline (Fig. 4A,B), but the modulation of the profile of the panorama was also reflected in the neural response of the TL neuron (arrow in Fig. 4B). To test whether the TL neurons combine both stimuli in a linear manner, as shown for the celestial cues, we also tested the same neurons’ responses to the single stimuli. As expected, the TL neurons responded to the sun stimulus and the panoramic scene, when presented individually (Fig. 4C, upper panel). Again, we used the neural tuning to the single stimuli to model the expected neural response of the TL neurons to a combined – sun and panorama – stimulus presentation. We then used the modeled neural response based on the weighted linear model (red curve in Fig. 4C) and correlated it with the actual neural response to both stimuli (yellow curve in Fig. 4C). In contrast to the results for the sun and polarized light (Fig. 2), the weighted linear model did not always result in high correlation values with the actual responses (Fig. 4D, inset, 0.55 ± 0.29, n = 13). Five of the 13 TL neurons (correlation coefficient < 0.5; Fig. 4D, inset) seemed to combine the sun stimulus and the panoramic skyline in a non-linear manner. The predicted weighting factors were highly variable. While some neurons exhibited responses that could be mainly explained by a response to the sun stimulus (weight < 0.5; Fig. 4D), responses of other TL neurons were more dominated by the simulated panoramic skyline (weight > 0.5; Fig. 4D). Taken together, we found a high variance in the coding of the cue preference between the sun and panoramic skyline information which stands in contrast to the results observed with the sun stimulus and the polarized light. Although we were not able to define from which TL subtype we obtained our recordings (see discussion), the results indicate that the cue hierarchy between celestial and terrestrial cues shows a high inter-individual flexibility in monarch butterflies.

**Fig. 4:**
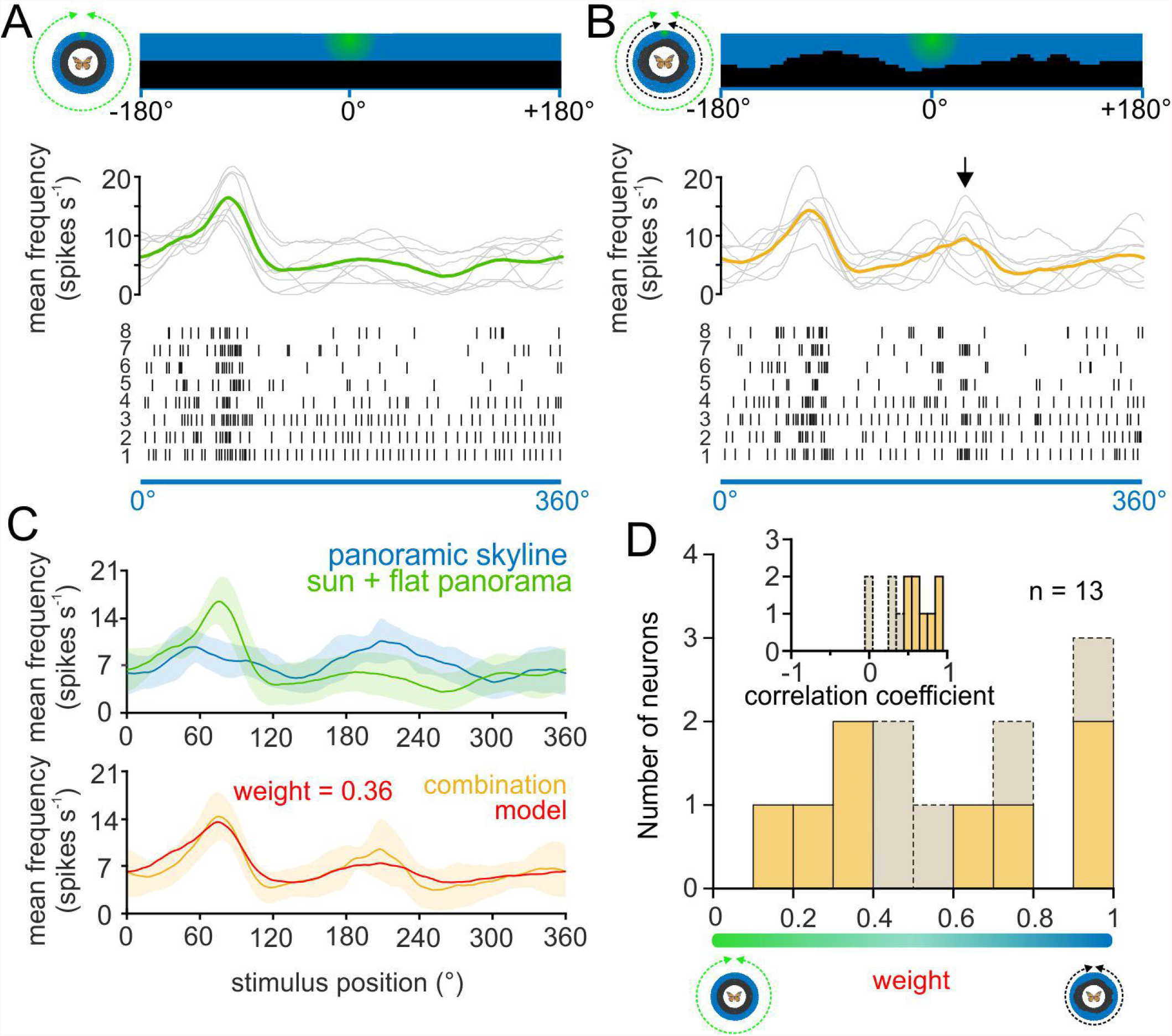
Weighting the sun and panoramic skyline in TL neurons. **(A,B)** The response of a TL neuron to eight 360°-rotations of the sun stimulus (A), and when the sun stimulus and the panoramic scene were rotated simultaneously around the animal (B). The stimulus position prior to rotation is shown (top). The action potentials rates are shown as raster plots (bottom). Above the raster plots, the sliding averages of the rotations are shown (middle plot, grey curves). The sliding averages of the mean spiking frequency are color coded (A, moving sun stimulus in green; B moving sun stimulus and panorama in yellow). Arrow indicates an increased spiking activity that can be contributed to the panoramic skyline **(C)** Upper plot: Sliding averages of the same TL neuron as in A and B responding to a 360° rotation of the panoramic skyline alone (blue curve) or the sun stimulus (green curve). Lower plot: sliding average of the activity of the same neuron when both stimuli (sun stimulus and panoramic skyline) were rotated simultaneously (yellow curve). A weighted linear model (red curve) was fitted to the observed neural activity to the combined stimulus. Shaded areas show the standard deviation. **(D)** The distribution of the weighting factors for experiments with the sun stimulus and panoramic skyline (n=13). Low weighting indicates that the sun stimulus and high weighting that the panorama stimulus dominated the combined response. The correlation coefficient distribution in the inset indicates how well the weighted linear model described the recorded neural response to the combined stimulus. Weighting factors obtained from a low correlation coefficient (<0.5), indicating that the neural response did not follow a linear model, are shown in grey.

## Discussion

We show that the monarch butterfly CX integrates multiple visual cues, i.e., celestial and terrestrial panoramic skyline cues for orientation. While the sun stimulus and polarized light were integrated linearly, the coding of the sun stimulus and the panoramic skyline did not always match a linear summation of action potential rates. Moreover, while polarized light was usually weighted stronger than the green sun stimulus, the weighting of the sun versus the panorama stimulus was set in a variable manner across different TL neurons. This observation is in line with behavioral results on monarch butterflies tested within the same visual setting and might allow the butterflies to set the cue preference in a highly flexible manner between celestial and terrestrial cues (Franzke et al., 2020).

### Celestial coding in the central complex

Monarch butterfly TL neurons are sensitive to polarized light (Heinze and Reppert, 2011; Nguyen et al. 2021, this work), similar to what has been reported for TL cells of a wide range of other insects, including desert locusts (Vitzthum et al., 2002, Heinze et al., 2009, Bockhorst et al., 2015, Pegel et al., 2018; Takahashi et al. 2022), field crickets (Sakura et al., 2008), dung beetles (el Jundi et al., 2015), sweat bees (Stone et al., 2019), and fruit flies (Hardcastle et al., 2021). In addition to polarized light, TL neurons in the present study were tuned to a green light spot – likely representing the sun – similar to what has been shown in TL-neurons of desert locusts (Pegel et al., 2018; Takahashi et al. 2022) and dung beetles (el Jundi et al., 2015), as well as in previous experiments in monarch butterflies (Heinze and Reppert, 2011; Nguyen et al., 2021). Thus, the TL neuron sensitivity to celestial cues is highly conserved and may play a crucial role for the heading coding in insects.

Importantly, we found a fixed spatial relationship between the sun stimulus and polarized light evoking the strongest neural responses. Like in Heinze and Reppert (2011), the here recorded TL neurons’ sensitivity to the simulated sun was strongly biased to the right side of the butterflies. In addition, the preferred firing direction of the same TL neurons to polarized light was significantly aligned with the longitudinal axis of the butterflies which is at odds with a previous study (Nguyen et al., 2021). This results in an orthogonal relationship of the mean sun stimulus and polarized light preferred directions, which parallels the 90°-relationship between the sun and polarization pattern in nature, as reported for TL neurons (Pegel et al., 2018) and optic lobe neurons (el Jundi et al., 2011) in the desert locust. However, Pegel et al. (2019) showed that the preferred directions to the sun stimulus differ between the three TL subtypes (TL2a,TL2b,TL3) in desert locusts. The monarch butterfly TL neurons can also be divided anatomically into three subtypes that innervate different layers in the lower division of the central body (Heinze et al., 2013). Unfortunately, we were not able to define from which subtype we performed our recordings as we often co-stained several TL subtypes in one experiment. As shown previously, monarch butterfly compass neurons show the same preferred firing direction, irrespective of the spectral information of the light stimulus (Heinze and Reppert, 2011; Nguyen et al., 2021). However, recordings from compass neurons in the desert locust suggest that the spectral influence on the preferred firing direction is strongly sensitive to the light intensity of the stimuli (Kinoshita et al., 2007; Pfeiffer and Homberg, 2007). It is therefore crucial to study the response characteristics of TL neurons to spectral cues at different light intensities in the future to shed light on how the monarch butterfly compass network may integrate different celestial cues into the central complex and how this represents the celestial cue hierarchy exhibited behaviorally.

### Integration of the panoramic skyline in the central complex

In previous experiments, the sensitivity of TL cells has been studied with respect to vertical stripes (Sun et al., 2017; Bockhorst and Homberg, 2017; Fisher et al., 2019), grating patterns (Rosner et al., 2019) or small light spots (Seelig and Jayaraman, 2013) in insects. We here found that TL neurons were sensitive to a simulated panoramic skyline scene by responding to changes in brightness while the panorama was rotated around the animal. Rather than encoding the current heading, these neurons seem to convey visual information into the insect compass, similar to the *Drosophila* ring neurons (Seelig and Jayaraman, 2013). In both monarch butterflies and fruit flies, they synapse on a population of neurons termed CL1 neurons (Heinze et al., 2013), called EP-G cells in fruit flies, which likely represent a distinct heading direction within a visual scene based on multimodal information (Seelig and Jayaraman, 2015; Kim et al., 2019; Turner-Evans et al., 2020; Beetz et al., 2021). How monarch butterfly CL1 cells compute a heading based on terrestrial information from TL cells awaits to be answered through neural recordings during flight as the coding strongly depends on the animal’s locomotory state (Beetz et al., 2021).

Most TL neurons seem to encode the panoramic skyline at elevations between -10° and 10° where the height of the panorama was most strongly modulated. This observation fits to behavioral results of Australian desert ants that mainly use the lower sector of a visual scenery for orientation (Graham and Cheng, 2009). However, as we only tested the response of the monarch TL neurons to one distinct panoramic scene, it has yet to be identified how modifying the frequency and amplitude of the panorama’s profile will affect the tuning of the TL neurons. We chose this specific panoramic skyline as a recent behavioral study showed that monarch butterflies are able to use this setting to sustain a directed flight course (Franzke et al., 2020). However, as their orientation performance was indistinguishable from a flight stabilization strategy, it was unclear whether the butterflies can employ compass orientation with respect to a panoramic scene. Our data here show that the central complex receives visual information of the panoramic scene, which suggest that monarch butterflies can use a panoramic skyline as a compass cue to compute a heading direction with respect to it.

### Flexible weighting between celestial and terrestrial information

When we presented the sun and polarization stimulus simultaneously to the butterflies, the TL neurons combined these cues in a linear manner. These results differ from the dung beetle TL neurons (el Jundi et al., 2015) but are in line with desert locust columnar CX-neurons (Pegel et al., 2019). The polarization UV stimulus was consistently ranked higher than the green sun stimulus in TL neurons in monarch butterflies when presented with a similar relative light intensity. When we presented a UV light spot instead of a green one, the unpolarized light stimulus dominated the neural tuning. This switch in cue preference was likely not a result of a change in wavelength but rather a consequence of a change in relative intensity of light, which is in line with the stronger response of TL neurons to UV light than to green light (Nguyen et al., 2021). Thus, as the sun is several magnitudes brighter than the remaining sky in nature, it is likely the dominant cue being encoded in TL neurons under a real sky.

In general, the weighting between the simulated sun and polarized light was very similar across different TL neurons. This low variability in cue preference between TL neurons recorded in different monarch butterflies was similar to what has been found for TL neurons in dung beetles (el Jundi et al., 2015) and might suggest that the weighting of celestial cues is determined at an early processing stage in the brain, such as at the level of the photoreceptors. In contrast, the simulated sun and the panoramic skyline were not always linearly integrated in the monarch butterfly central complex. Moreover, the cue preference was highly variable, which is well in line with the high interindividual difference in the behavioral use of these cues for orientation in a flight simulator (Franzke et al., 2020). This high flexibility indicates that the weighting might not only be set based on the sensitivity of the inputs at the butterfly’s eye but might additionally be adjusted at later stages in the brain network. This would allow a high inter-individual difference in weighting that is based on the animal’s internal state, as well as its experience.

## Author contributions

Study design: TATN, CM, KP, BeJ. Conducting experiments: TATN. Analysis of data: TATN. Interpretation of data: TATN, MJB, CM, KP, BeJ. Drafting of the manuscript: TATN, BeJ. Critical review of the manuscript: MJB, CM, KP. Acquired Funding: BeJ. All authors approved of the final version of the manuscript.

## Acknowledgments

We thank Dr. James Foster for fruitful comments on our analysis and Kolja Richter and Konrad Öchsner for their help in developing the LED arena. We thank Samantha Iiams, Aldrin Lugena, Guijun Wan and Ying Zhang for their help in capturing monarch butterflies and for checking them for *Ophryocystis elektroscirrha*. In addition, we would like to thank Sergio Siles and Marie Gerlinde Blaese (butterflyfarm.co.cr) for providing us with monarch butterfly pupae.

